# IGF2BP3 Promotes Autophagy-Mediated TNBC Metastasis via M6A-Dependent, Cap-Independent c-Met Translation

**DOI:** 10.1101/2024.09.23.614652

**Authors:** Ziwen Wang, Yihan Li, Mengyuan Cai, Xu Zhang, Ruoxi Xu, Haiyan Yang, Yuzhou Huang, Liang Shi, Jifu Wei, Qiang Ding

## Abstract

Tumor metastasis involves a series of complex processes and is a major challenge in clinical treatment. The cooperation of epigenetic modifications and metabolic adaptations endows tumor cells with dynamic adaptations for survival in variable environments, which is crucial for tumor metastasis and worth exploring in depth. In this study, we found IGF2BP3 could link epigenetic modification and metabolic adaptation in promoting autophagy-mediated triple-negative breast cancer (TNBC) metastasis. As a TNBC specifically high expressed N6-methyladenosine (m6A) binding protein, IGF2BP3 could bind to the m6A motif of c-Met mRNA, regulating autophagy-mediated epithelial-to-mesenchymal transition (EMT) progress by c-Met/PI3K/AKT/mTOR pathway. Moreover, through the recruitment of eIF4G2 as a collaborator, IGF2BP3 induced the changes in c-Met protein expression not by affecting mRNA stability, but by facilitating its mRNA translation initiation in an m6A-dependent and cap-independent manner. In conclusion, our study extended the role of IGF2BP3 in autophagy mediated TNBC metastasis, found IGF2BP3 could bind to the m6A motif on the 5′, 3′-UTR of c-Met mRNA to facilitate its translation in a cap-independent manner.

## Introduction

Triple negative breast cancer (TNBC), which characterized by the absence of estrogen receptor, progesterone receptor, and human epidermal growth factor receptor 2 expression (Bianchini, Balko, Mayer, Sanders, & Gianni, 2016), is the most aggressive subtype of breast cancer (Nolan, Lindeman, & Visvader, 2023). It shows a lower overall survival rate and higher relapse rate, compared to other subtypes (Bianchini, De Angelis, Licata, & Gianni, 2022). Although TNBC is highly responsive to systemic chemotherapy, the easily metastatic properties make it an early tendency to metastasize with a high recurrence rate, and a poor prognosis. Moreover, after recurrence and metastasis, TNBC usually progresses faster and shows strong resistance to chemotherapy and radiotherapy, which is a major pain point in its clinical treatment (Bai, Ni, Beretov, Graham, & Li, 2021). Therefore, clarifying the intrinsic mechanism of the high metastatic nature in TNBC gives the chance to acquire its precision treatment.

Except as a cytosolic component required for organelles recycling and intracellular pathogens clearance, autophagy is increasingly recognized for its important role in cancer metastasis (Su, Yang, Xu, Chen, & Yu, 2015). The relationship between autophagy and epithelial-to-mesenchymal transition (EMT) is one of the main characters. EMT introduces epithelial cells with adheren junctions, desmosomes, and tight junctions morphed into stromal cells which are loosely distributed in the stroma, giving tumor cell abilities to migrate (Thiery, Acloque, Huang, & Nieto, 2009). The occurrence of the EMT process is always accompanied by the autophagy activation due to their similar related signaling pathways activation (H.-T. Chen et al., 2019). Autophagy inhibitors and activators like chloroquine, 3-methyladenine, rapamycin have demonstrated therapeutic effects in multiple cancers including TNBC by regulating EMT in clinical and preclinical experiments (Abd El-Aziz, Gillson, Jansson, & Sahni, 2022; Singla & Bhattacharyya, 2017; So et al., 2016; H. Wang, Zhang, Wu, Wang, & Wang, 2018). However, it is widely accepted that the role of autophagy in EMT is bidirectional: autophagy activation provides energy and metabolites which are harmful substances to EMT during the advanced tumor metastasis and help cells survive in stressful environments. On the flip side, autophagy downregulates key transcription factors of EMT in early tumor stages and thereby impede metastasis (Gundamaraju et al., 2022). Its complexity adds the difficulty to Figure out the specific regulatory processes, hindering the development and clinical use of related drugs. Therefore, further explorations of its mechanisms in TNBC metastasis deserve more attention.

As the most common post-transcriptional modification of eukaryotic RNAs (Zhao, Roundtree, & He, 2017), N6-methyladenosine (m6A) influences various processes of cancer development (Huang, Weng, & Chen, 2020; X. Lin et al., 2019; Liu et al., 2022; Nag, Goswami, Das Mandal, & Ray, 2022). Up to now, its regulators such as methyltransferase3 (METTL3), FTO alpha-ketoglutarate dependent dioxygenase and YTH n6-methyladenosine RNA binding protein C1 (YTHDC1) have been reported to occupy an important position in cancer autophagy by regulating related RNAs splicing, degradation, and translation (X. Chen et al., 2022; Hao et al., 2022; He et al., 2022; Liang et al., 2022; Ning et al., 2023; Yang et al., 2022). In our previous study, we found insulin-like growth factor 2 mRNA-binding protein 3 (IGF2BP3), an m6A reading protein, was specifically expressed in TNBC and introduced RNA-seq, RIP-seq, MeRIP-seq to reveal its function (X. Zhang et al., 2023). After the secondary analysis, RIP-seq hinted IGF2BP3’s significance in cell migration and autophagy at the same time. This phenomenon hinted us that IGF2BP3 may alter cell migration capacity by affecting cellular autophagy and gave us a chance to further interpret the relationship between cellular autophagy and TNBC metastasis at the molecular level. IGF2BP3 may identify the m6A motifs of related RNAs, promoting their stability by recruiting co-factors like ELAV-like RNA-binding protein 1, matrin 3, and polyadenylate-binding protein 1 (Zhu, Hong, & Ling, 2023). These regulations may lead to RNA abundance changes. But many targets with unchanged RNA levels are neglected and the regulatory mechanism is still unclear. Previous study suggested IGF2BPs might change translation efficiency, but only found its location change between ribosome and non-ribosome fractions and this phenomenon happened in heat shock situation (Huang et al., 2018). Inspired by previous study that eukaryotic translation initiation factor 3 (eIF3) may recognize m6A motif and improve translation initiation in a cap-independent manner (Meyer et al., 2015), we conjectured whether IGF2BP3 could regulate RNA through the translation initiation without altering its abundance.

In this study, we discovered that IGF2BP3 played a crucial role in TNBC cell autophagy, leading to TNBC metastasis. Furthermore, we tried to elucidate the underlying molecular mechanism of this phenomenon in TNBC and finally given a viewpoint that IGF2BP3 could target the m6A motif on 5′, 3′-untranslated regions (UTR) of cellular-mesenchymal to epithelial transition factor (c-Met) mRNA and increase its mRNA translation in a cap-independent manner by recruiting eukaryotic translation initiation factor 4 gamma 2 (eIF4G2) to affect autophagy mediated TNBC metastasis.

## Result

### Downregulation of IGF2BP3 could enhance the autophagy in TNBC cells

Based on our previous study, KEGG gene enrichment analysis with RNA immunoprecipitation (RIP)-seq were joined to highlight IGF2BP3’s role in autophagy and cell adhension through binding transcripts (Figure 1A). To verify this, except MDA-MB-231 cell line used in previous study (X. Zhang et al., 2023), BT-549 cells were also transfected with lentivirus to overexpress or suppress IGF2BP3 expression, qRT-PCR and western blot were used to validate it (Figure 1-figure supplement 1A, B). The overexpression of IGF2BP3 significantly increased the autophagy receptor protein P62 expression, and hindered the autophagy marker LC3 transformed from inactive (LC3-Ⅰ) to activated (LC3-Ⅱ) form at the same time in TNBC cells (Figure 1B). Overexpression of IGF2BP3 showed the contrary tendency (Figure 1C). Subsequently, autophagosome and autophagolysosome were also observed visually by transmission electron microscopy, suggesting that decreased IGF2BP3 could upregulate the autophagy in TNBC cells (Figure 1D). Finally, we utilized the GFP-mCherry-LC3 reporter to observe the dynamic changes from initial autophagosome formation to autophagolysosome formation. Since lysosomes maintained a stable acidic circumstances (Hu, Chen, Liu, & Xu, 2023) and GFP fluorescence quenched easily in acidic environment, the presence of red dots indicated just mCherry fluorescence when autophagosomes phagocytosed by lysosomes, forming autophagolysosomes. Yellow dots, on the other hand, indicated the overlap of GFP and mCherry fluorescence, signifying initial autophagosomes. After transfection with shIGF2BP3 lentivirus, both yellow and red dots increased briefly, especially the yellow dots (Figure 1E, F). This phenomenon indicated that IGF2BP3 might be involved in the regulation of initial autophagosome formation. All the results provided compelling evidences that downregulation of IGF2BP3 could enhance the autophagy in TNBC cells.

**Figure 1:**
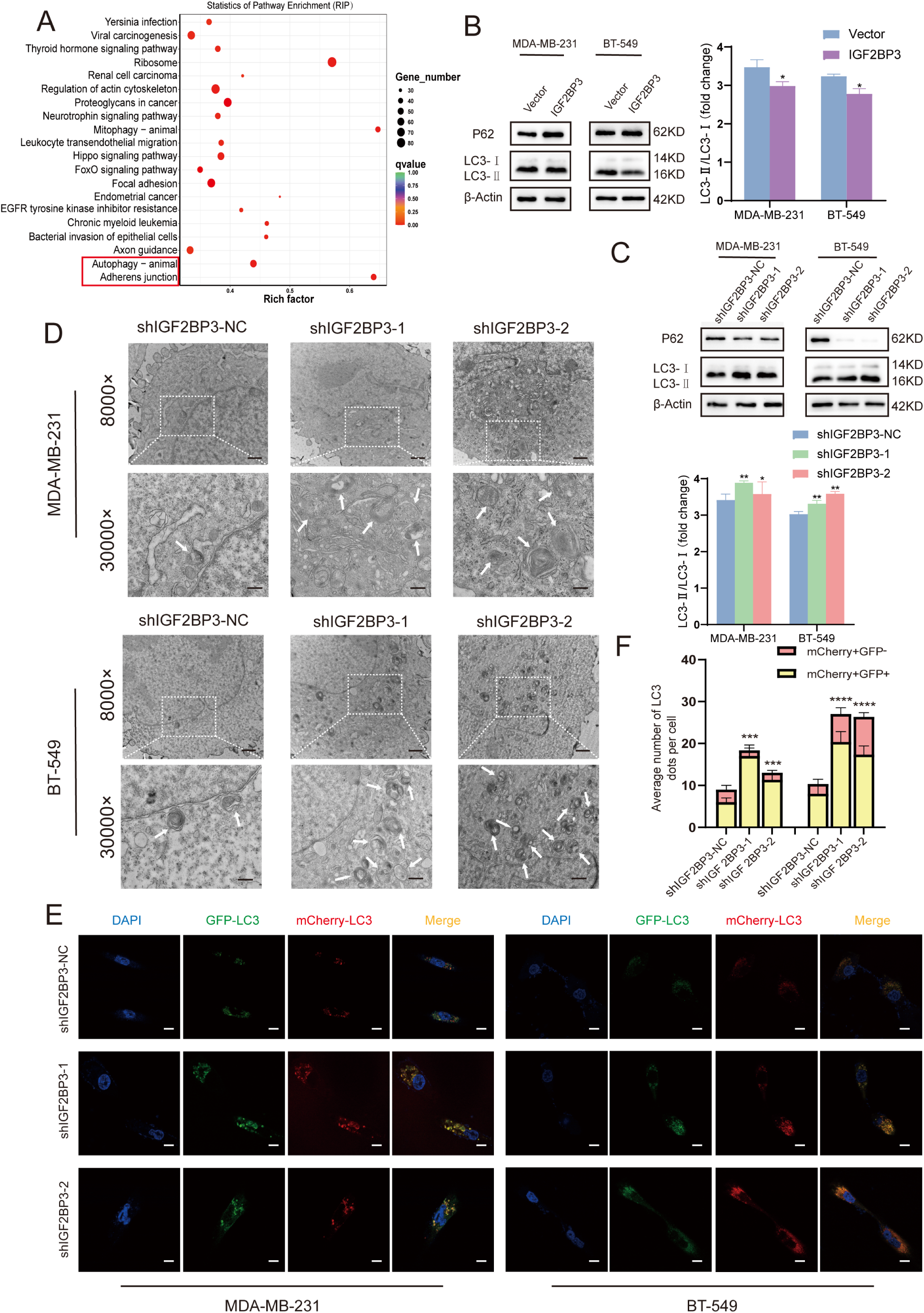
downregulation of IGF2BP3 could enhance the autophagy in TNBC cells. A. KEGG pathway analyses were identified to show the enrichment in RIP-seq of IGF2BP3. Autophagy and cell adhension pathway were in the top 10. B. Western blot assays exhibited P62, LC3 expression after the overexpression of IGF2BP3 in MDA-MB-231 and BT-549 cells. The gray value of LC3-Ⅱ /LC3-Ⅰ representing the activation of autophagy. C. Western blot assays exhibited P62, LC3 expression after the knockdown of IGF2BP3 in MDA-MB-231 and BT-549. The gray value of LC3-Ⅱ /LC3-Ⅰ representing the activation of autophagy. D. Electron micrographs demonstrated the fine structure of autophagosomes or autophagolysosomes in MDA-MB-231 and BT-549 cells after the knockdown of IGF2BP3. Scale bar, 1 μm; Scale bar in the magnified image, 0.5 μm. E-F. GFP-mCherry-LC3B reporter and IGF2BP3 shRNA were transfected in MDA-MB-231 and BT-549 cells to show the autophagy flux changed by IGF2BP3. Changes in yellow and red fluorescence were observed by confocal microscope. Red and yellow dots indicated autophagolysosomes and autophagosomes, respectively. Scale bar, 10 μm. Data were shown as mean ± SEM, *P <0.05 (one-way ANOVA).

**Figure 1-figure supplement 1:**
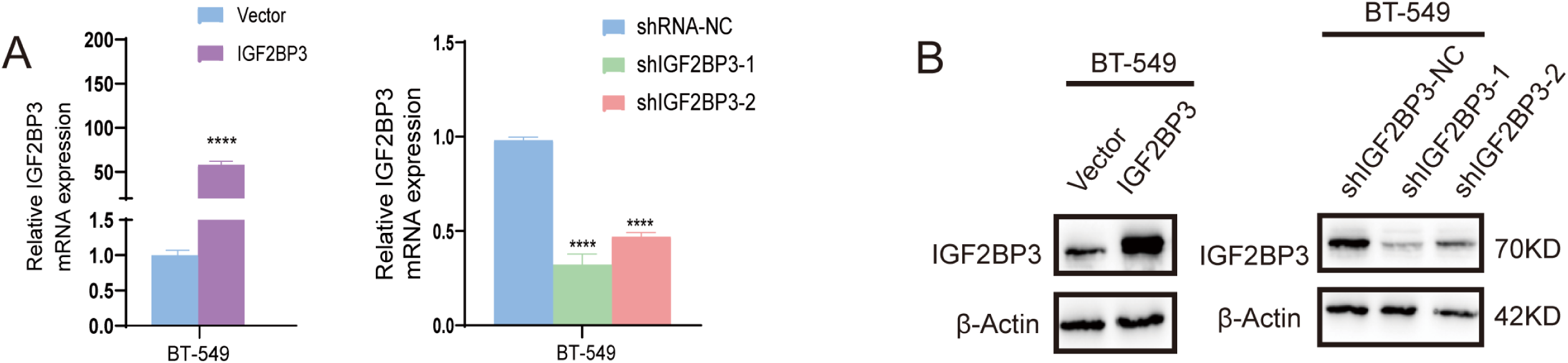
transfected efficiency of IGF2BP3 in BT-549 cell. A-B. BT-549 cells were transfected with lentivirus to overexpress and knockdown IGF2BP3. Transfection efficiencies were verified by qRT-PCR (A) and western blot (B).

### IGF2BP3 hindered cell autophagy to promote the EMT of TNBC cell

To test whether the inhibition of cell autophagy by IGF2BP3 could affect TNBC metastasis, we firstly conducted wound healing and transwell assays to verify the migratory capabilities of TNBC cells. The overexpressed IGF2BP3 led to wound healing area (Figure 2-figure supplement 1A, B) and cell shuttle numbers (Figure 2-figure supplement 1 C-E) increase in TNBC cells. Likewise, knocking down IGF2BP3 resulted in a substantial decrease in TNBC cells migration under the same conditions (Figure 2-figure supplement 1 A-E). Then, we examined the epithelial phenotype marker E-cadherin and mesenchymal phenotype markers, N-cadherin, Vimentin by western blot. E-cadherin was up regulated, N-cadherin and Vimentin were down regulated in IGF2BP3 knockdown groups and opposite trends were exhibited in IGF2BP3 overexpression groups (Figure 2-figure supplement 1F). Subsequently, mTORC1 inhibitor rapamycin which treated as a classical autophagy agonist was introduced into both the control and shIGF2BP3 TNBC cells. P62, LC3 expression (Figure 2-figure supplement 2A) and GFP-mCherry-LC3 reporter (Figure 2-figure supplement 2B, C) showed the stimulative effect of autophagy. After rapamycin added, wound healing (Figure 2A, C) and transwell assays (Figure 2B, D) showed the activation of autophagy weakened the TNBC cells migration ability and reversed the changes induced by IGF2BP3 alteration partially. At the same time, the upregulation of E-cadherin and the downregulation of N-cadherin, Vimentin due to the reduction of IGF2BP3 were hindered in TNBC cells (Figure 2E). In conclusion, IGF2BP3 could accelerate the EMT process by inhibiting cell autophagy, thereby promoting TNBC cell migration.

**Figure 2:**
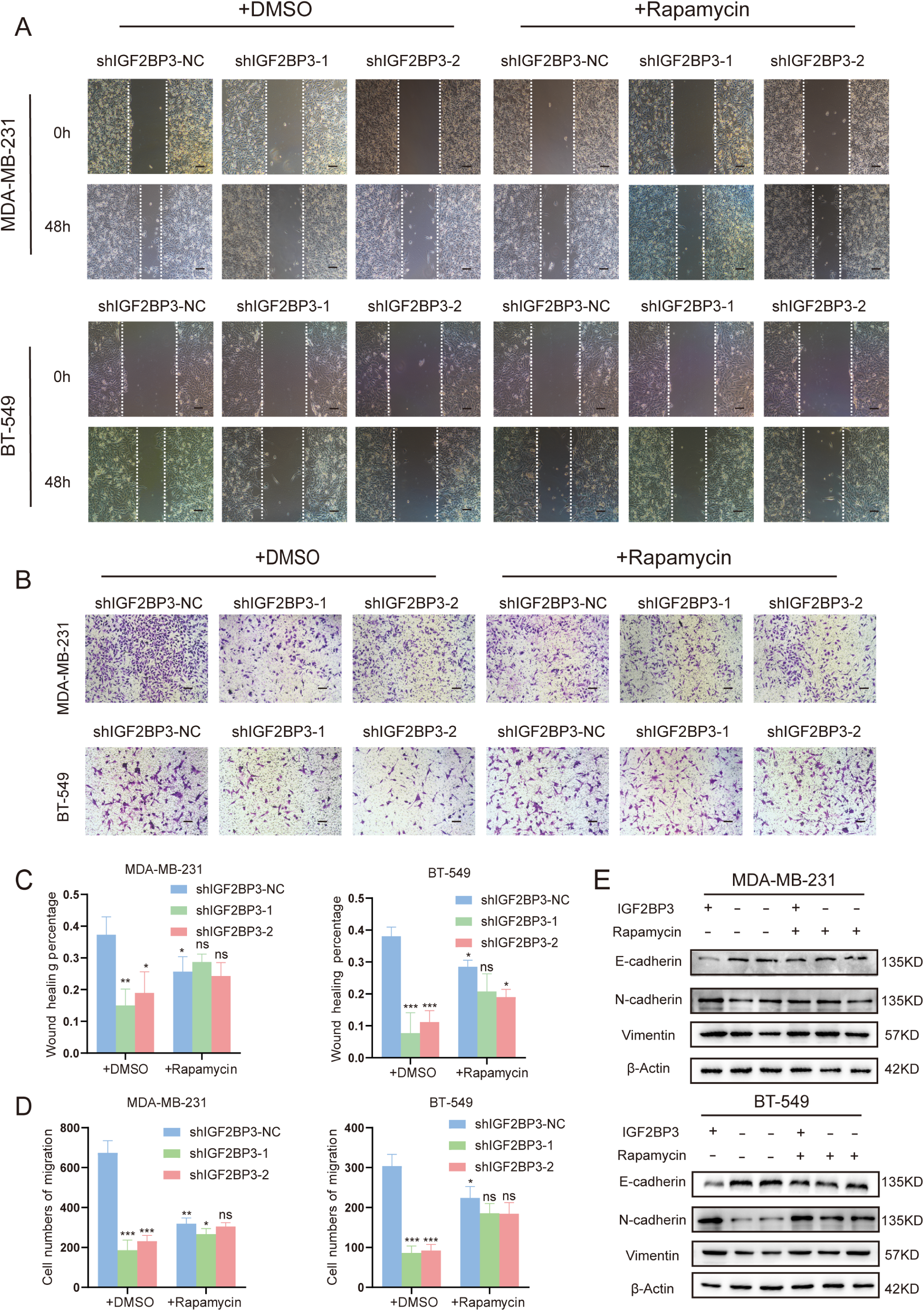
TNBC metastasis affected by IGF2BP3 was reversed after the activation of autophagy. A, C. Wound healing assay was performed to detect MDA-MB-231 and BT-549 cells migration level after the knockdown of IGF2BP3 and joined rapamycin treatment (10 μM, 48 h) (original magnification, 10×) (bars, 50 μm). Student t test was used for the comparison between groups. Data were shown as the mean ± SEM of three replicates; *P < 0.05. B, D. Migration ability of MDA-MB-231 and BT-549 cells changed by the knockdown of IGF2BP3 and joined rapamycin treatment were exhibited by transwell assay. Scale bar, 50 μm. E. The expression of E-cadherin, N-cadherin and Vimentin in MDA-MB-231 and BT-549 cells was detected by western blot after the knockdown of IGF2BP3 and joined rapamycin treatment.

**Figure 2-figure supplement 1:**
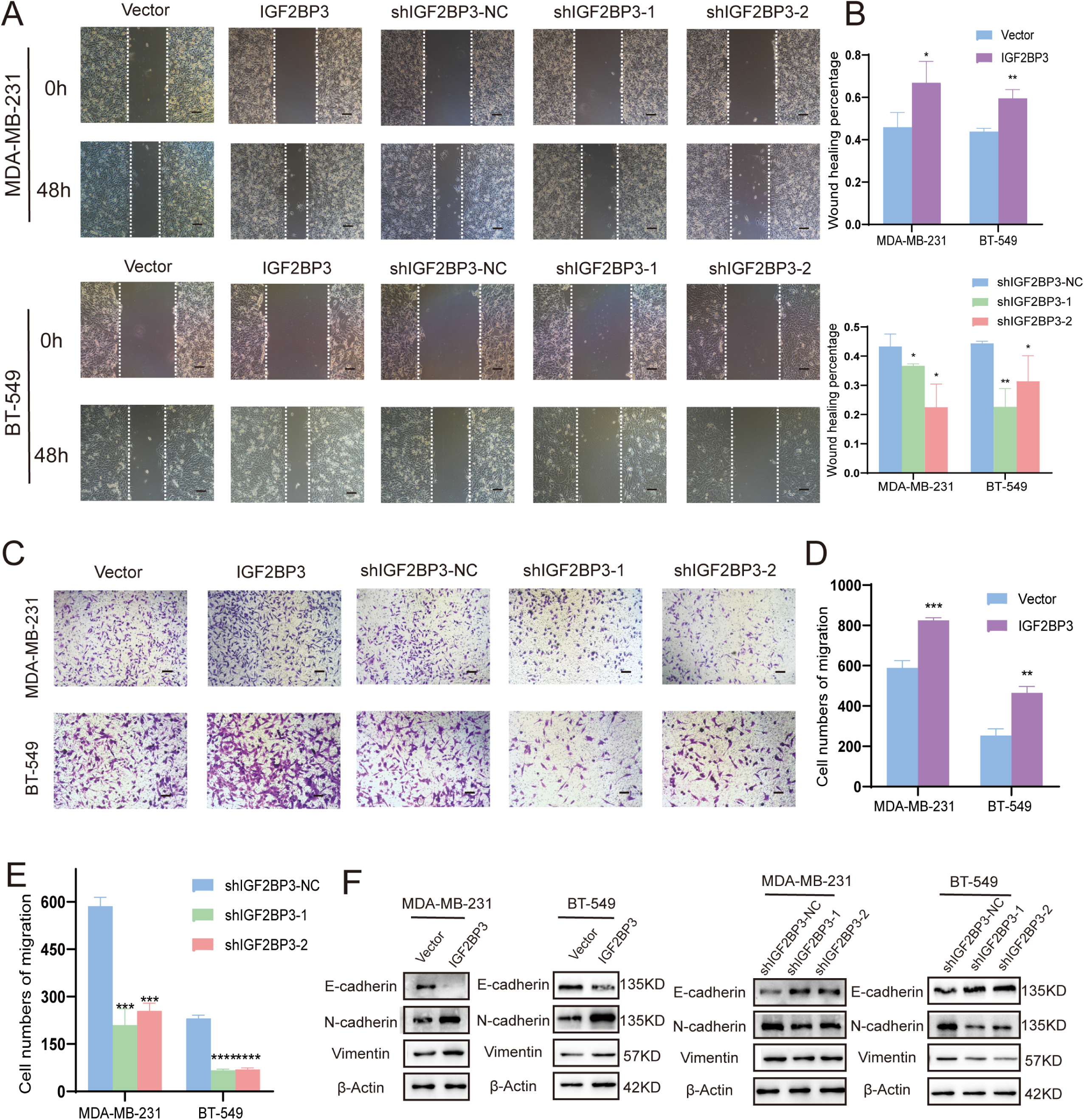
IGF2BP3 promoted TNBC cell migration through EMT progression. A-E The migration level in MDA-MB-231 and BT-549 cells after knockdown of IGF2BP3 were tested by wound healing assay (A, B) and transwell assay (C-E). F. The expression of epithelial phenotype marker E-cadherin, mesenchymal phenotype markers N-cadherin and Vimentin in MDA-MB-231 and BT-549 cells was detected by Western blot after overexpression or knockdown of IGF2BP3.

**Figure 2-figure supplement 2:**
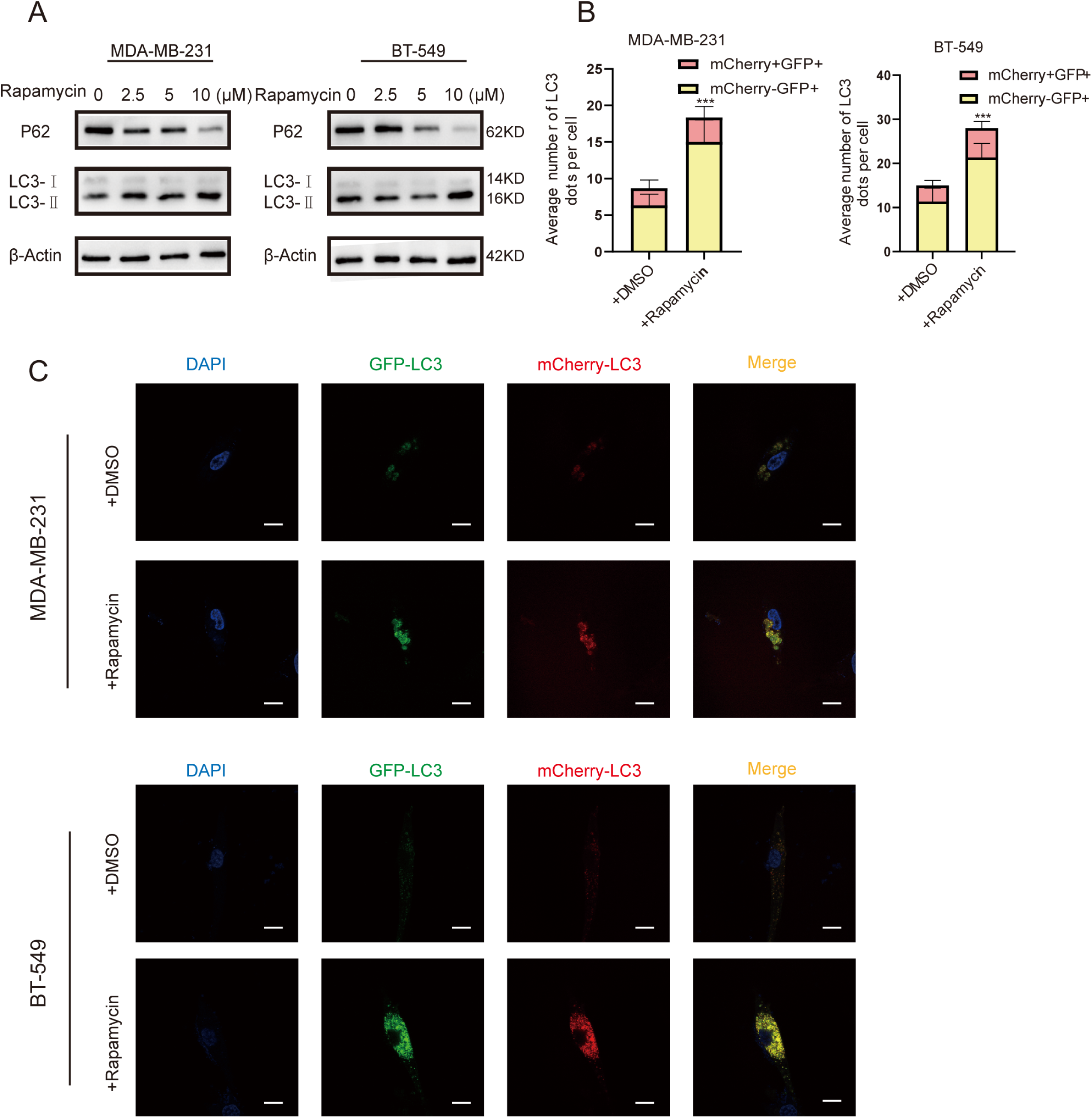
rapamycin activated TNBC cell autophagy. A. The expression of LC3 and P62 after rapamycin treatment in MDA-MB-231 and BT-549 cells were shown by western blot. B-C. Autophagosomes and autophagolysosomes changed in MDA-MB-231 and BT-549 cells after rapamycin treatment were by confocal microscopy. MDA-MB-231 and BT-549 cells were transfected with GFP-mCherry-LC3B reporter. Red and yellow dots indicated autophagolysosomes and autophagosomes, respectively.

### c-Met could be the target for IGF2BP3 functioned in autophagy mediated TNBC cell migration

Excepted RIP-seq, RNA-seq and MeRIP-seq were also introduced to employed a comprehensive analysis. After overlapping, 699 genes bound by IGF2BP3 were marked with m6A, of which only 129 genes exhibited altered RNA expression upon the knockdown of IGF2BP3, while the majority of them (570) remained unaffected (Figure 3A). However, it’s worth noting that previous study predominantly focused on the targets of IGF2BP3 exist RNA alterations (Z. Lin et al., 2023; Xin Wang et al., 2021; N. Zhang et al., 2022). So, we brought in KEGG pathway cluster to analyze and classify the transcripts (570) (Figure 3B). At the same time, we screened out highly expressed genes in TNBC from the TCGA database (Figure 3-figure supplement 1A) and united them with the genes from autophagy inhibitory pathway: phosphoinositide 3-kinase (PI3K)/ protein kinase B (AKT)/ mechanistic target of rapamycin (mTOR) and EMT related adheren junction pathway. Among these, c-Met emerged as the sole candidate meeting these criteria (Figure 3C). To clarify the correlation between IGF2BP3 and c-Met in *vitro*, we collected 16 tissue specimens of TNBC from our institution and qRT-PCR demonstrated their high correlation of expression (Figure 6D). Moreover, high expression of c-Met implied poor prognosis in breast cancer patients demonstrated from Kaplan-Meier Plotter database (Figure 6E). RIP-qPCR (Figure 3D) and the binding peak positions (Figure 3E) cleared the direct co-binding of IGF2BP3 protein with c-Met mRNA. Human breast cancer samples from TCGA database (Figure 3G) and various human breast cancer cell lines (Figure 3-figure supplement 1B, C) proved c-Met mRNA’s specific high expression in TNBC. qRT-PCR and mRNA stability analysis validated that c-Met’s mRNA abundance (Figure 3H) and half-life period (Figure 3-figure supplement 2A, B) didn’t change obviously followed by overexpression or knockdown of IGF2BP3. On the contrary, western blot showed that c-Met’s protein expression represented significant changes with the alteration of IGF2BP3 and demonstrated a high degree of protein correlation with IGF2BP3 in TNBC patient samples from CPTAC database (Figure 3F). At the same time, c-Met’s downstream PI3K-AKT-mTOR’s phosphorylation level was also been influenced (Figure 3I). To Figure out whether IGF2BP3’s impact on c-Met protein expression depend on promoting protein synthesis or inhibiting protein degradation, cycloheximide (CHX) was added to block protein synthesis. Finally, no significant changes were observed in c-Met protein degradation rates following the knockdown of IGF2BP3 (Figure 3J). This suggests that IGF2BP3 may primarily affect protein synthesis, thereby regulating c-Met protein expression. Based on the above results, c-Met was identified to be a direct and unique target of IGF2BP3 which functioned in autophagy mediated migration in TNBC, mainly modulating c-Met expression at the protein synthesis process.

**Figure 3:**
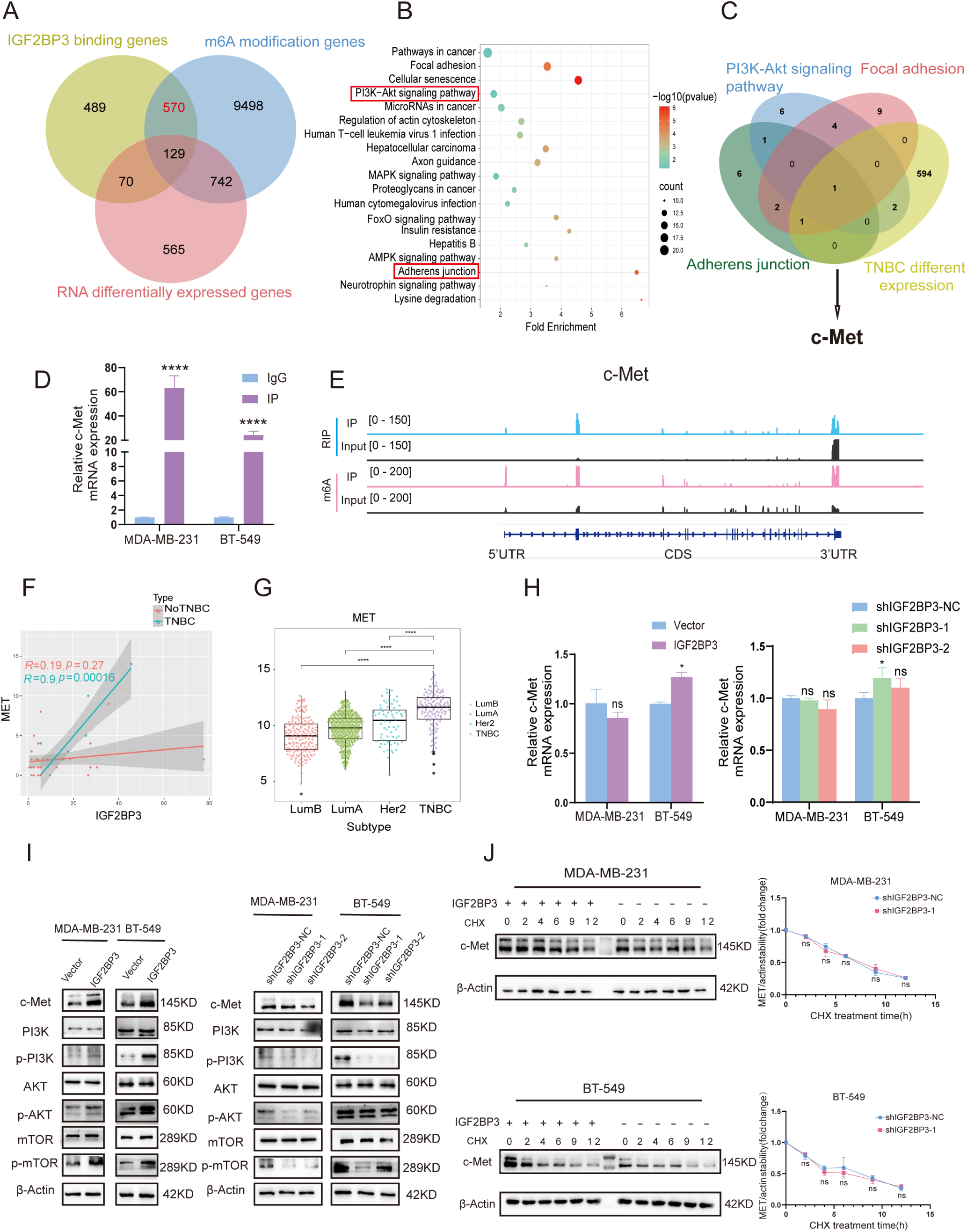
IGF2BP3 could affect autophagy mediated TNBC metastasis by increasing c-Met protein expression. A. Venn diagram showed the intersection genes of RIP-seq, m6A-RIP-seq, and RNA-seq. B. KEGG pathway analysis of genes intersected by RIP-seq and m6A-RIP-seq. C. Overlapping analysis of genes identified by PI3K-AKT signaling pathway/ adherens junction/ focal adhesion/ TNBC different expression genes. D.RIP-qRT-PCR illuminated the direct connection between IGF2BP3 and c-Met mRNA. E. Distribution of IGF2BP3-binding peaks and m6A peaks across c-Met mRNA. F. Correlation analysis between IGF2BP3 and c-Met’s protein expression in no TNBC patient samples (n = 35) and TNBC patient samples (n = 11) from CPTAC database. High degree of protein correlation between IGF2BP3 and c-Met was found in TNBC patient samples G. mRNA expression of c-Met in different subtype breast cancer from TCGA database. H. mRNA expression of c-Met after overexpression or knockdown of IGF2BP3 determined by qRT-PCR. I. Western blot showed c-Met protein and PI3K-AKT-mTOR phosphorylation levels changed by IGF2BP3. J. Control or shIGF2BP3 MDA-MB-231 and BT-549 cells were treated by cycloheximide to inhibit protein synthesis. The c-Met protein degradation efficiency was demonstrated by western blot.

**Figure 3-figure supplement 1:**
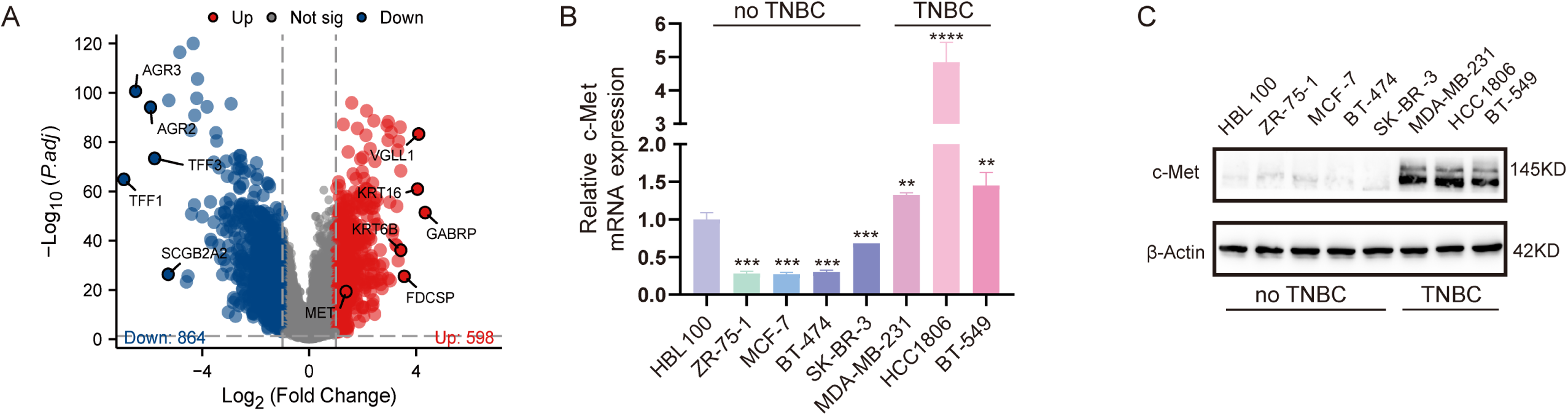
c-Met is high expressed in TNBC. A. Volcano plot shown differentially expressed genes in TNBC tissues compared with non-TNBC tissues. Data from TCGA database. B-C: mRNA and protein expression of c-Met in different subtype breast cancer cell were detected by qRT-PCR and western blot.

**Figure 3-figure supplement 2:**
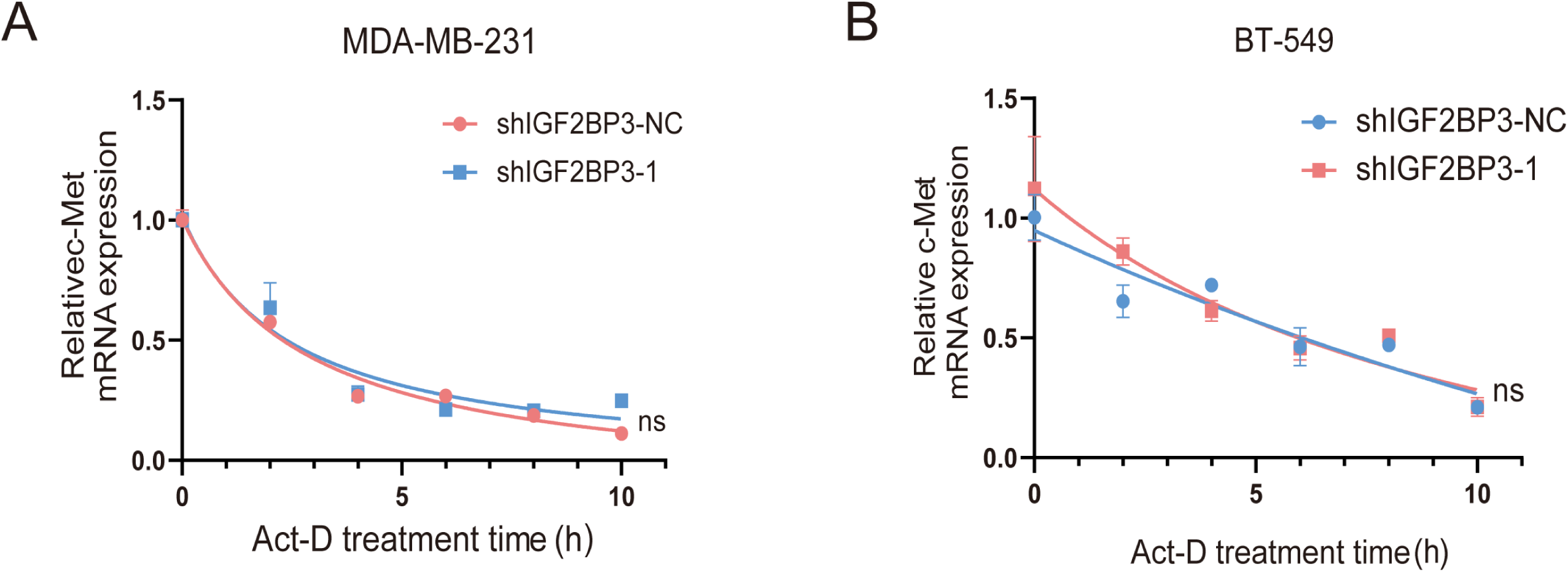
IGF2BP3 has no concern with c-Met mRNA stability. A-B. MDA-MB-231 and BT549 cells which transfected with shIGF2BP3 were treated with 5 μg/mL actinomycin D (ActD) for 0, 2, 4, 6, 8, 10 h, c-Met mRNA stability was calculated by qRT-PCR.

### IGF2BP3 bound to the m6A motifs on 5′, 3′-UTR of c-Met mRNA

As illustrated in RIP-seq, IGF2BP3 binding peaks were distributed among exon, transcription start site, intron and 5′, 3′-UTR regions (Figure 4A). Additionally, focusing binding peaks on c-Met revealed by RIP-seq, most of the IGF2BP3 binding sites scattered among the 5′-UTR, CDS and 3′-UTR regions of c-Met mRNA, closely aligned with the m6A peaks (Figure 3E). Therefore, we initially selected the m6A site of c-Met mRNA as the binding site of IGF2BP3 for validation. Given the canonical m6A base motif been "RRACH" [32], we identified sequences rich in “RRACH” within the 5′-UTR, CDS and 3′-UTR regions of c-Met mRNA (each spanning approximately 500 nt), and constructed 3 dual luciferase reporter plasmids based on these sequences (Figure 4C). We transformed them into TNBC cells separately and found the luciferase activity of the cells carrying 5′-UTR or 3′-UTR base sequences plasmid dropped when IGF2BP3 was knockdown (Figure 4D). Then, we introduced point mutations at the potential m6A loci on the 5′-UTR and 3′-UTR regions of c-Met mRNA, substituting “A” with “T” and transfected plasmids same as before (Figure 4E). Consistent with our expectations, the alterations observed in the wild type (WT) groups vanished in the mutation (MUT) groups (Figure 4F). These results confirmed that IGF2BP3 could bind to c-Met mRNA in an m6A-dependent manner. To further narrow this scope, transcripts were fragmentized to approximately 100 nt segments and primers were designed to specifically amplify the fragments containing the above m6A mutation sites. MeRIP-qPCR confirmed the direct m6A modification on these mRNA fragments (Figure 4G) and biotin RNA probes were designed accordingly (Figure 4H). RNA pull-down assay using these RNA probes successfully captured IGF2BP3 proteins (Figure 4I). Thus far, we have established that IGF2BP3 could bind randomly to m6A sites within the 5’ and 3’-UTR regions of c-Met mRNA, narrowing down the binding region to 70-100 nucleotides.

**Figure 4:**
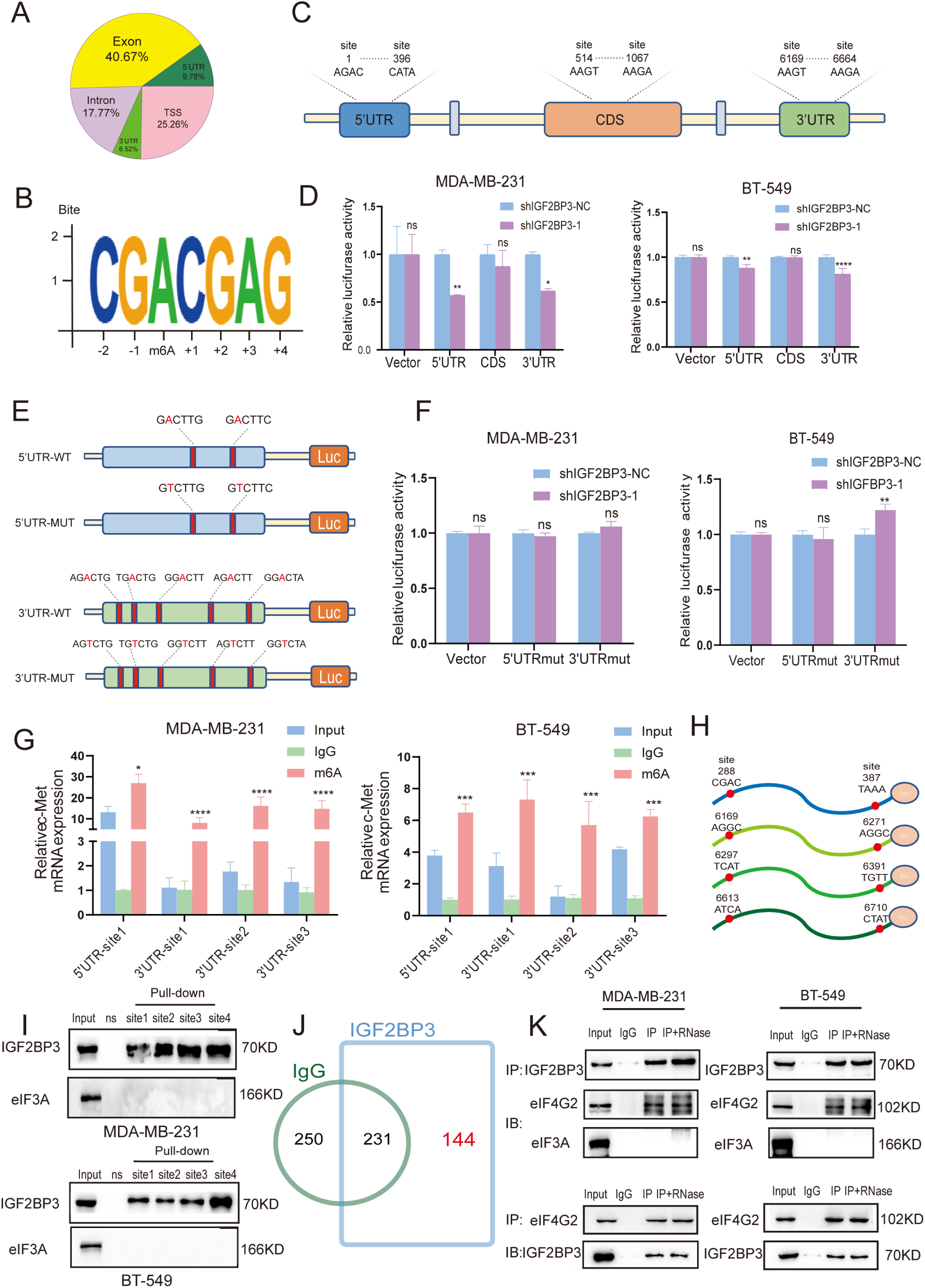
IGF2BP3 interacted with eIF4G2 in promoting c-Met mRNA translation through an m6A-dependent manner. A. IGF2BP3 binding peak distribution on transcripts showed by RIP-seq. B. m6A enrichment motif detected by the Homer motif analysis with m6A-seq data. C-D. Control and shIGF2BP3 MDA-MB-231 and BT-549 cells were transfected with dual luciferase plasmid containing 5′-UTR, CDS or 3′-UTR base sequences of c-Met mRNA (C), the luciferase activity changed by IGF2BP3 was tested by the dual-luciferase reporter assay system (D). E. Mutant m6A motif in c-Met mRNA 5′-UTR, 3′-UTR, changed “A” to “T”. F. Transfected the m6A site mutation (MUT) sequences dual luciferase plasmid in control and shIGF2BP3 MDA-MB-231 and BT-549 cells and luciferase activity were showed. G. MeRIP-qRT-PCR verified the direct m6A modification in 5′-UTR and3′-UTR of c-Met mRNA. H. RNA biotin probes for m6A motif in 5′-UTR and 3′-UTR of c-Met mRNA were designed and proven in (G). I. Pull-down assay was conducted using biotin probes to exhibit the binding of IGF2BP3 or eIF3A at the m6A modification site of c-Met mRNA. J. Proteins co-immunoprecipitated with IGF2BP3 detected by LC-MS. K. Western blot showed the co-immunoprecipitating relationship between IGF2BP3, eIF4G2 and eIF3A.

### IGF2BP3 interacted with eIF4G2 to stimulate c-Met translation in an m6A-dependent manner

Previous findings suggested that m6A in 5′ UTR of mRNAs could recruit eIF3 to promote cap-independent mRNA translation and m6A regulators such as YTHDF1/2/3 binding at 5′-UTR and 3′-UTR of mRNAs could cooperate with eIF3 in promoting mRNA translation (S. Lin, Choe, Du, Triboulet, & Gregory, 2016; Meyer et al., 2015; A. Wang et al., 2023; Xiao Wang et al., 2015). So, we hypothesized that IGF2BP3 might cooperate with eIF3 to influence mRNA translation progression as YTHDF1/2/3. Unfortunately, RNA probes failed to capture eIF3A proteins while successfully catching IGF2BP3 proteins (Figure 4I) and IGF2BP3 couldn’t combine with eIF3A by co-IP (Figure 4K). In this phenomenon, the immunoprecipitation products induced by IGF2BP3 were sent for mass spectrometry identification. After excluding non-specific binding to IgG, a total of 144 proteins were detected to have protein-protein interactions with IGF2BP3 (Figure 4J). Inside them, eIF4G2 was picked out as a member of eIF4 complex, mediated cap-independent translation initiation (Liberman et al., 2015). Western blot validated the interaction and the addition of ribonucleases revealed that the interaction between them was RNA-independent (Figure 4K). Combining with the above results, we identified IGF2BP3 and eIF4G2 had mutual effect thus regulating c-Met mRNA translation initiation in an m6A-dependent, cap-independent manner.

### c-Met played a pivotal role in mediating the effects of IGF2BP3 on autophagy progression in TNBC

To substantiate that the function of IGF2BP3 in autophagy-mediated metastasis of TNBC was generated through c-Met, we overexpressed c-Met in both control and shIGF2BP3 TNBC cells. mRNA and protein levels of c-Met were detected by qRT-PCR and western blot (Figure 5-figure supplement 1A, B). Notably, the GFP-mCherry-LC3 reporter also represented a reduced number of autophagosomes and autophagolysosomes after transfection with the c-Met plasmid in both control and shIGF2BP3 groups (Figure 5A, B, Figure 5-figure supplement 1C). At the same time, c-Met overexpression resulted in decreased P62 levels and increased LC3 expression in comparison to the NC group. Also, when suppressing IGF2BP3 and overexpressing c-Met at the same time, changes caused by IGF2BP3 suppression were reversed by c-Met overexpression (Figure 5C, D). Consistently, transwell assay (Figure 5E, F), wound healing assay (Figure 5-figure supplement 1D, E) and western blot (Figure 5C, D) showed that both cell migration ability and EMT progress suppressed by IGF2BP3 suppression were re-established after c-Met overexpression. Furthermore, the phosphorylation level of PI3K-AKT-mTOR pathway increased in tandem with c-Met overexpression (Figure 5G, H). Taken together, our data demonstrated that c-Met played a pivotal role in mediating the effects of IGF2BP3 on autophagy progression in TNBC, consequently enhancing its metastasis.

**Figure 5:**
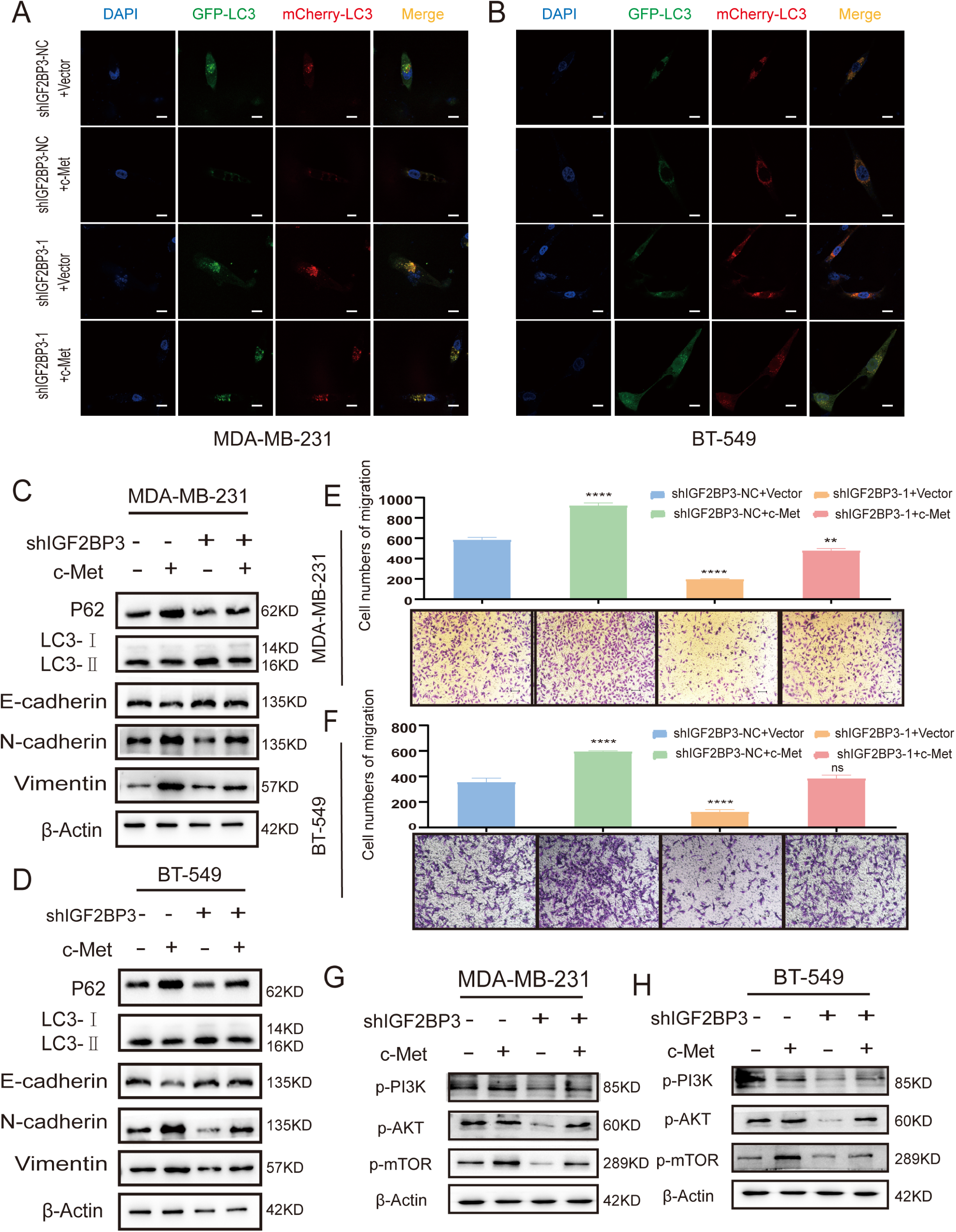
IGF2BP3 promoted autophagy mediated TNBC metastasis depending on c-Met. A-B. Autophagy flux changes after overexpression of c-Met and knockdown of IGF2BP3. Yellow and red dots indicated autophagosomes and autolysosomes. Scale bar, 10 μm. C-D. EMT-related markers and autophagy-related markers altered by overexpression of c-Met. E-F. The number of cell migration was changed after knockdown of IGF2BP3 and overexpression of c-Met was demonstrated by transwell assay. Data were shown as mean ± SEM, *P <0.05 (Student t test). G-H. PI3K-AKT-mTOR phosphorylation expression after knockdown of IGF2BP3 and overexpression of c-Met were demonstrated via western blot.

**Figure 5-figure supplement 1:**
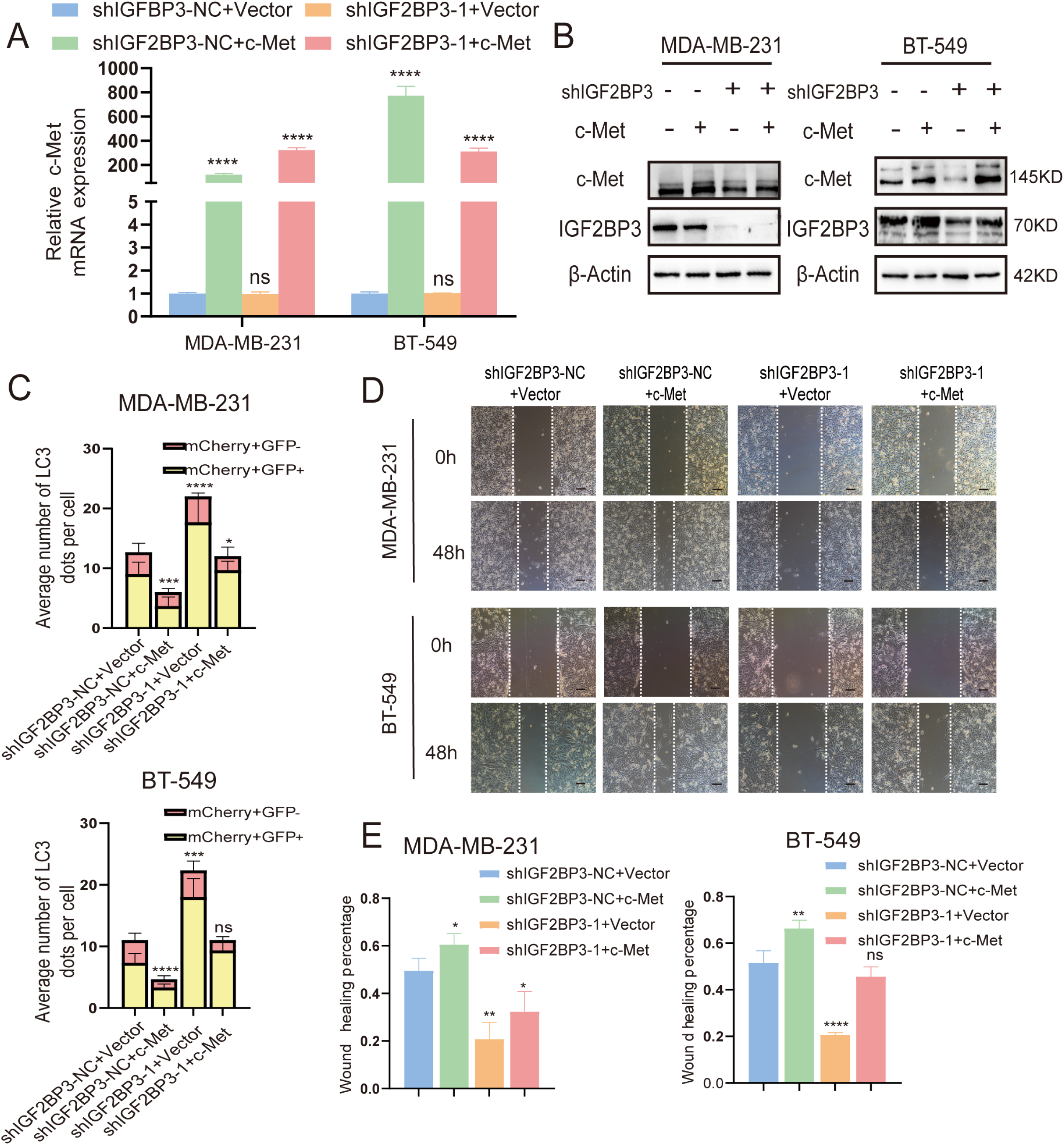
altered cellular autophagy and migration capacity resulting from knockdown of IGF2BP3 was rewritten by c-Met overexpression. A-B. Negative control and shIGF2BP3 group of MDA-MB-231 and BT-549 cells were transfected to overexpress c-Met, mRNA and protein level verified by qRT-PCR and western blot. C. Autophagy flux activated by the knockdown of IGF2BP3 were inhibited by the overexpression of c-Met in MDA-MB-231 and BT-549 cells. Data were shown as mean ± SEM, *P <0.05 (one-way ANOVA). D-E. Cell migration ability suppressed by IGF2BP3 suppression were re-established after c-Met overexpression in MDA-MB-231 and BT-549 cells.

### IGF2BP3 relied on c-Met for enhancing tumor metastasis in breast cancer lung metastasis mice model

To verify the function of IGF2BP3 in *vitro* having reproducibility *in vivo,* shIGF2BP3-NC+Vector, shIGF2BP3-NC+c-Met, shIGF2BP3-1+Vector, and shIGF2BP3-1+c-Met transfected MDA-MB-231 cells were utilized to establish a breast cancer lung metastasis mice model. Cells were injected into nude mice via tail veins and mice lung tissue samples were collected at day 35. The lung tissue sections were further analyzed by H&E staining (Figure 6A) and total numbers of lung metastases were counted. Remarkably, the overexpression of c-Met significantly induced an increase in the number of lung metastases in mice, and the inhibition of IGF2BP3 resulted in a decreased number. Notably, the changes observed in the latter group were reversed partially by the former (Figure 6C). Additionally, immunohistochemical results revealed that IGF2BP3 and c-Met level were positively correlated with N-cadherin level and negatively correlated with LC3 level (Figure 6B). To sum up, in accordance with the trend in *vitro*, IGF2BP3 and c-Met collectively improved the TNBC metastasis and transform autophagy marker LC3 and EMT maker N-cadherin protein level in breast cancer lung metastasis model. Taken together, our data demonstrated that IGF2BP3 relies on c-Met for enhancing tumor metastasis in mice breast cancer lung metastasis model.

**Figure 6:**
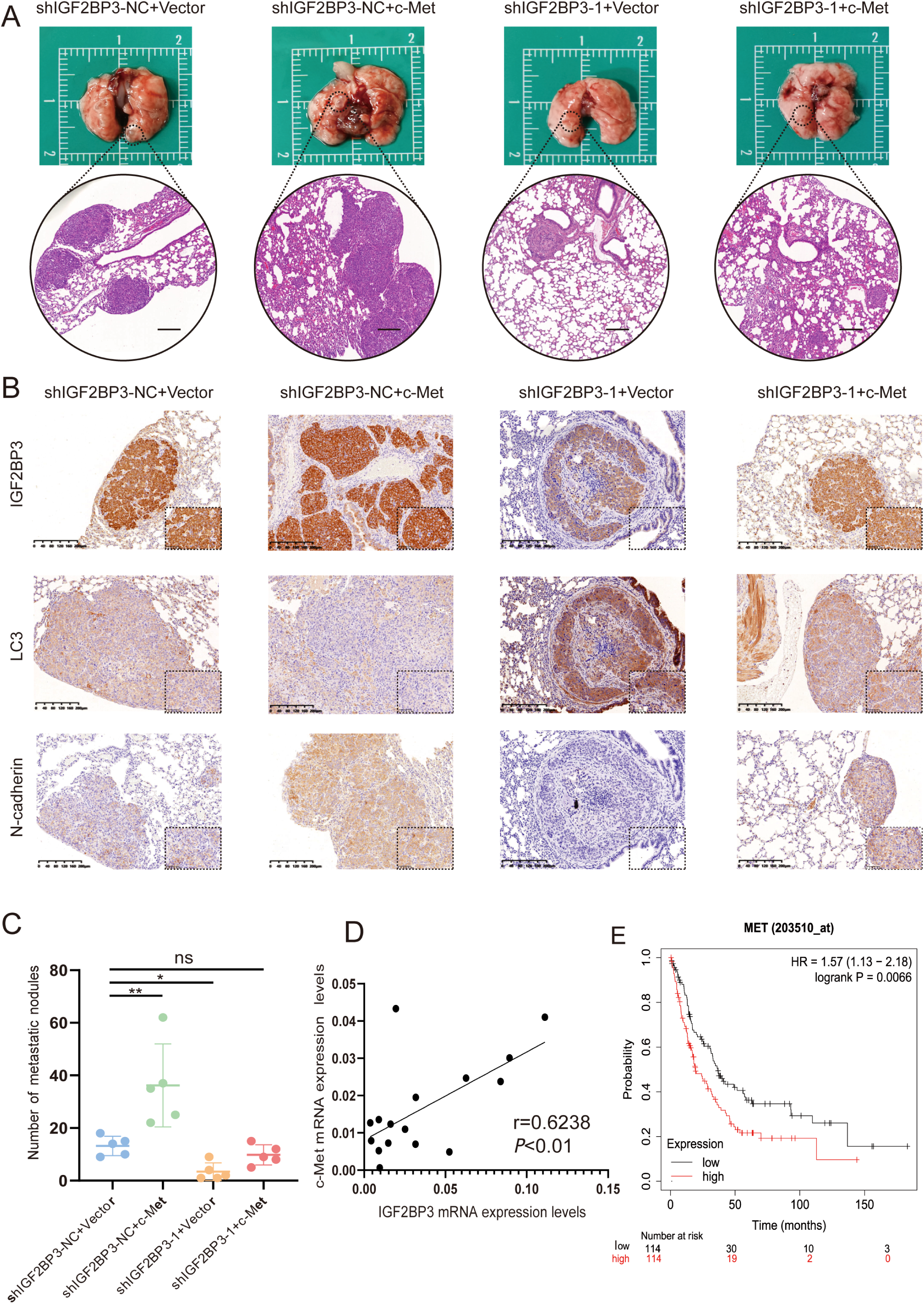
IGF2BP3 relied on c-Met for enhancing tumor metastasis in *vivo*. A. Representative images revealed the lung tissue metastasized by MDA-MB-231 cells which overexpressed c-Met or knockdown IGF2BP3. HE staining of lung section (n=5) showed the pathological properties of the nodules. Scale bar, 400 μm. B. The expression of IGF2BP3, LC3, N-cadherin were analyzed by immunohistochemistry in paraffin-embedded tissue (original magnification, 20×) Scale bar, 400 μm. C. The number of breast cancer lung metastasis nodules of the four groups were calculated. Data were shown as mean ±SEM, \**P*<0.05. D. qRT-PCR demonstrated high correlation of expression between IGF2BP3 and c-Met in TNBC patient tissues (n = 16). E. Kaplan–Meier analysis showed high expression of c-Met implied poor prognosis in breast cancer patients from Kaplan-Meier Plotter database.

**Figure 7:**
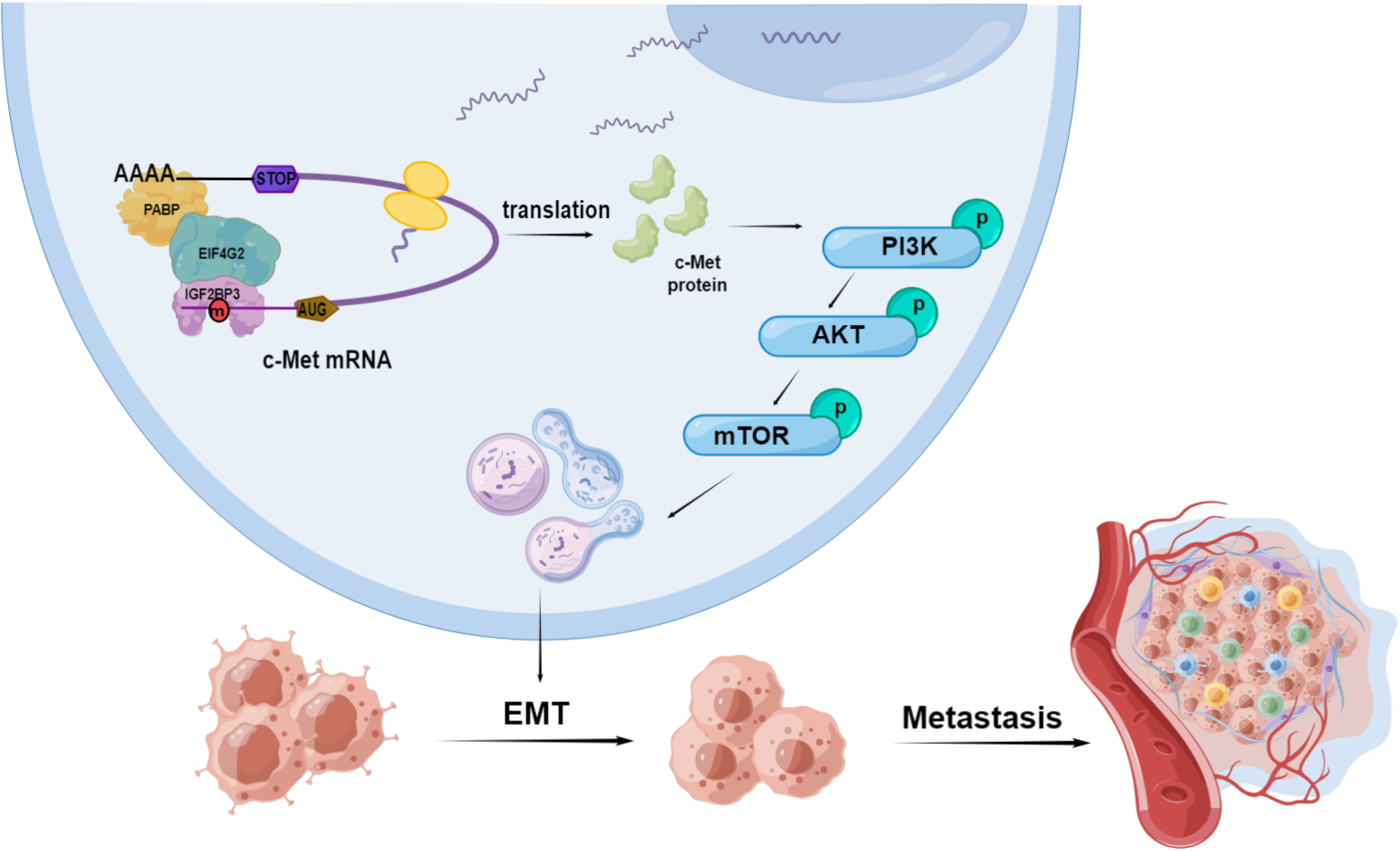
a proposed model of IGF2BP3 joined in autophagy mediated TNBC metastasis with m6A-dependent molecular mechanisms. IGF2BP3 interacts with eIF4G2 to enhance c-Met translation in an m6A-dependent manner. c-Met promoted autophagy-mediated metastasis of TNBC via phosphorylating PI3K-AKT-mTOR pathway.

## Discussion

In previous study, IGF2BP3 functioning as an m6A “reader” played a pivotal role in regulating RNA translation, stability, degradation and splicing (Zhu et al., 2023). We discovered its specific high expression in TNBC and joined in cell proliferation (X. Zhang et al., 2023), but its function is far from being fully elucidated. In the present study, we uncovered that IGF2BP3 could inhibit the autophagy in TNBC cells, thereby promoting the transition of cells from an epithelial to a mesenchymal phenotype, which further facilitated TNBC metastasis.

Tumor metastasis involved a series of biological events. The primary tumor cells acquire the ability to invade the basement membrane and deeper tissues through self-remodeling, and then spread in the circulation and effectively evade the killing of immune cells. Eventually, they colonize distant organs and resume proliferation (Gerstberger, Jiang, & Ganesh, 2023). These events depend on the strong dynamic adaptability of tumor cells to survive in the variable environment. However, the purely genetic mechanism struggles to cope with these changes in the short term, making dynamic epigenetic and metabolic adaptations dominant (Massagué & Ganesh, 2021). These two factors complement each other and jointly reshape highly metastatic tumor cells. As an essential m6A “reader”, IGF2BP3’s role in breast cancer metastasis has been reported. Kim *et al*. found IGF2BP2 and IGF2BP3 promoted TNBC metastasis cooperatively by destabilizing progesterone receptor (H.-Y. Kim, Ha Thi, & Hong, 2018). Also, Jing *et al*. verified that IGF2BP3 might repress slit guidance ligand 2 expression in facilitating TNBC metastasis (Jiang et al., 2022). These studies gave some explanations on mechanisms by which IGF2BP3 promoted metastasis, but ignored the co-interaction of metabolic adaptations in it. Based on RIP-seq, we found transcripts which combined with IGF2BP3 clustered highly in autophagy and cell migration pathway at the same time. This hints us that IGF2BP3 may endow TNBC cell with a stronger metastatic property in the mediation of autophagy as an active metabolic process. As we knew, autophagy specifically phagocytoses intracellular proteins, organelles, and metabolites by the formation of double-membrane vesicles. Through the key proteins and metabolites phagocytosis, autophagy may involve in central biological processes of cell migration like focal adhesion dynamics, integrin signaling transport, EMT and tumor stromal cell interaction (Dower, Wills, Frisch, & Wang, 2018). Based on our experiments, we found IGF2BP3 could inhibit TNBC cell autophagy and promote its migration, and autophagy activator rapamycin was used in clarifying the role of autophagy in this process. As a specific mTORC1 inhibitor, rapamycin formed a complex with FKBP12 and gave this complex the ability to allosteric mTORC1(Li, Kim, & Blenis, 2014). The allosteric mTORC1 lose their kinase activity, failed to inhibit the ULK1/2 and the VPS34 complex phosphorylation, and the complex after phosphorylation induced autophagosome membrane extension, thus autophagy initiation, leading to the activation of autophagy by rapamycin (Y. C. Kim & Guan, 2015). After treatment with rapamycin, we found EMT progression in TNBC was impeded. Meanwhile, cell migration changes of TNBC caused by knockdown of IGF2BP3 was reversed partially. These results suggested that IGF2BP3 could regulate TNBC metastasis process by autophagy-mediated EMT.

As a momentous receptor tyrosine kinase, c-Met is found to be mutated, amplificated and overexpressed in many cancers and thus promotes their progression by phosphorylating the signaling pathways like PI3K/AKT, JAK/STAT, Ras/MAPK and so on. The phosphorylation of these pathways always triggers cancer cells autophagy and EMT activation (Y. Zhang et al., 2018). Through our study, c-Met was found to be the target in autophagy-mediated EMT by IGF2BP3 in TNBC. In the previous studies, IGF2BP3’s role in post-transcriptional regulation were mostly focused on their RNAs stability alteration. However, IGF2BP3 was found to change the c-Met’s protein level and related phosphorylation regulatory activity, instead of its mRNA level in this study. More functions like RNA splicing or translation have been proposed and were worthy of in-depth investigation (An & Duan, 2022). After treatment of CHX to inhibit protein synthesis, c-Met protein degradation ratio didn’t decline by the knockdown of IGF2BP3. This result enlightened us that IGF2BP3 might change the c-Met expression by promoting protein synthesis.

Protein synthesis mainly refers to the translation initiation, elongation and termination, and occurs at different locations of mRNAs (Brito Querido, Díaz-López, & Ramakrishnan, 2024). In our study, we found the m6A binding sites of IGF2BP3 focused on the 5′, 3′-UTR of c-Met mRNA, which were also the translation initiation regions. At these regions, more than nine eukaryotic initiation factors function in start site identification, 43S initiation complex assembling and mRNA activation. Their combination and cooperation form a complete translation initiation process (Jackson, Hellen, & Pestova, 2010). However, these combinations are not unique and they bind to specific regions on the mRNA to form different modes of translation initiation such as cap-dependent, cap-independent, leaky scanning [47]. M6A regulators METTL3, YTHDF1/2/3 were reported to promote protein translation by cooperating with eIF3 in a cap-independent translation (Hao et al., 2022; S. Lin et al., 2016; A. Wang et al., 2023; Xiao Wang et al., 2015). EIF3 could recognize the m6A motif and recruit 43S initiation complex to initiate translation directly (Meyer et al., 2015). Interestingly, IGF2BP3 and eIF3 couldn’t cooperate with each other or target the same m6A site on c-Met mRNA in our study. We finally found eIF4G2 acted as a cooperator and helped IGF2BP3 in its mRNA translation. EIF4G2 is an emerging member in eIF4G family. Different from other family member initiating the translation in cap-dependent manner, eIF4G2 modulates cap-independent translation. It associates with eIF2β and eIF4AI and starts the translation from 5′-UTR internal ribosome entry site of mRNA (Liberman et al., 2015). Combing these results, we concluded that IGF2BP3 could recruit eIF4G2, instead of eIF3, to the m6A motif in 5′-UTR of c-Met mRNA and facilitate protein translation in a different cap-independent manner. In summary, our finding expounded a new impact of IGF2BP3 in TNBC. By targeting c-Met and improving its protein translation, IGF2BP3 achieved the function in TNBC metastasis via inhibited cell autophagy, not only highlighting the importance of the m6A methylation machinery in autophagy but also further confirming IGF2BP3’s multiple effect in TNBC. It confirmed the potential role as the therapeutic target in TNBC. Moreover, IGF2BP3 could bind to the m6A motif on 5′-UTR of its mRNA and motivate its protein translation by a novel cap-independent manner, extending its function and binding motif as an m6A reader.

## Materials&Methods

### Cell culture

Breast epithelial cell line HBL-100 and human TNBC cell lines: MDA-MB-231, BT549, HCC-1806; luminal breast cancer cell lines: MCF-7, ZR-75-1; HER2-breast cancer cell lines: BT474, SK-BR-3 were acquired from American Type Culture Collection (ATCC, USA). Cell lines were cultured in high glucose DMEM (Wisent, China) except HCC-1806 which was cultured in RPMI-1640 (Wisent, China). Both of them supplemented with 10% fetal bovine serum, 100 μg/ml penicillin-streptomycin (Hyclone, USA). The cells were incubated at 37°C in a cell culture incubator with 5% CO_2_.

### Lentivirus and plasmids transfection

Three different oligonucleotide sequences of lentiviral shRNAs and one full coding sequence (CDS) sequences of lentiviral plasmid synthesized by Obio Technology (Shanghai, China), were used to knockdown and overexpress IGF2BP3. In addition, GFP-mCherry-LC3 lentiviral was constructed by GenePharma (Shanghai, China). Finally, the stable cell lines were selected with 3 μg/ml puromycin. For overexpression of c-Met, CDS sequences of c-Met were subcloned into pcDNA3.1 and produced by Corues Biotechnology (Nanjing, China). Plasmids were transfected with Lipofectamine 3000 (Invitrogen). The sequences of lentivirus and plasmids were listed in appendix1-table1.

### RNA extraction and quantitative real-time PCR (qRT-PCR)

Total RNAs were extracted from cells by Trizol (Takara, Japan). HiScript Q RT SuperMix (Vazyme, China) was used to reverse the transcription of mRNAs. qRT-PCR was programed with the Roche LightCycler 480 RT-PCR system (Roche, Switzerland). The specific PCR primers were listed in appendix2-table1.

### Western blot

Cells were lysed in RIPA buffer (P0013C, Beyotime, China) with enzyme inhibitors. Protein concentration was determined by BCA protein assay kit (P0012, Beyotime, China). Sample was separated by SDS-PAGE and transferred to the PVDF membrane (Millipore, USA). The membranes were blocked with skim milk, incubated with relevant primary antibodies and HRP-conjugated secondary antibodies (CST, USA). The relative levels of targeted protein were normalized by β-actin and detected through Immobilob™ Western Chemiluminescent HRP Substrate (Millipore, USA). Primary antibodies joined in the experiments were listed in appendix3-table1.

### Transmission electron microscopy (TEM)

One million MDA-MB-231 and BT549 cells transfected with shIGF2BP3 lentivirus and negative control were gathered and fixed in 2.5% glutaraldehyde in phosphate buffer (0.1 M, pH 7.0) for 8 h, following 1% osmium tetroxide for 1 h. After that, a graded alcohol series dehydrated the samples and phosphotungstic acid negatively stained them. Finally, the ultrastructure was observed by electron microscope (JEOL JEM-1400Flash).

### GFP-mCherry-LC3 fluorescence imaging

Cells were transfected with GFP-mCherry-LC3 and shIGF2BP3 lentiviruses. After construction of stable strain, 1×10^4^ cells were inoculated in confocal dishes and immobilized by 4% paraformaldehyde. Nucleuses were dyed with 4’,6-diamidino-2-phenylindole. Finally, fluorescence photography was performed under a confocal microscope (Stellaris STED, Germany).

### Wound healing assay

After transfected IGF2BP3 overexpression and knockdown lentivirus, 1×10^6^ cells were seeded in a 6-well plate. Afterwards, 200 μl sterile pipette tips were used to create the linear scratch and washed twice with PBS to remove floating cells following to replace the media with serum free media. The wounded areas were photographed at 0 h and 48 h under the microscope. Image J was used to measure the area.

### Transwell assay

Tissue culture plate insert (Millicell Hanging Cell Culture Insert, USA) were implanted into 24-well plate. At the same time, 3×10^5^ cells were seeded on the top plate with serum-free DMEM, the bottom plates were filled with 600 μl medium with 10% FBS as a cell crossing catalyst. After 24 h or 48 h incubation, cells that successfully crossed were stained with 1% crystal violet.

### RNA immunoprecipitation (RIP)

Cells were cleaved with lysis buffer (Magna RIP Kit; Millipore, USA) and then incubated with 5 μg of anti-IGF2BP3 or anti-IgG at 4°C overnight. The RNA-protein complexes were captured by protein A/G magnetic beads. The collected RNAs were reversely transcripted and analyzed by qRT-PCR after RNA eluted and purification.

For methylated RNA immunoprecipitation (MeRIP), firstly, total RNAs were extracted from cells. Then, mRNAs were fragmented to 100 nt. After that, fragmented RNAs were hatched with m6A antibody and performed the following immunoprecipitation by MeRIP m6A kit (Millipore, Germany). The final products were also analyzed by qRT-PCR.

### Protein/RNA stability analysis

After transfected IGF2BP3 knockdown and control lentivirus, 5×10^5^ cells were plated into 6-well plates and then added into cyclohexane (CHX, 20 μg/ml) or actinomycin D (ActD, 5 μg/ml) at 0, 2, 4, 6, 9, 12 h. Total proteins and RNAs were extracted at the appropriate time. Then, c-Met protein and RNA degradation rate were detected by Western blot and qRT-PCR.

### Luciferase assay

After transfected IGF2BP3 knockdown and control lentivirus, cells were transfected with containing negative and particular c-Met sequence (c-Met-5′-UTR, c-Met-5′-UTR-mut, c-Met-CDS, c-Met-3′-UTR, c-Met-3′-UTR-mut) pGL3 reporter and luciferase vector. Luciferase activities were tested by the Dual-Luciferase reporter assay system (Promega, USA).

### Co-immunoprecipitation(co-IP)

The magnetic beads (MCE, USA) were incubated with IgG/IGF2BP3 anti-bodies at 4 °C for 2 h. At the same time, cells were lysed in 1ml IP buffer (P0013, Beyotime, China). Collected the supernatant co-incubated with magnetic bead antibody complex. After that, the beads were washed and separated from proteins. Total proteins sent to the Beijing genomics institute for liquid chromatograph-mass spectrometer (LC-MS/MS) and Western blot analysis was used to verify.

### RNA pull-down assay

Biotinylated RNA probes were synthesized by RiboBio (GenePharma, Shanghai, China) and captured with streptavidin-conjugated agarose magnetic beads (Smart-lifesciences, China). Then, cells were lysed and lysates were hatched with beads-RNA probe complex at 4 ℃ for 2 h. RNA Probe sequences joined in the experiment were listed in appendix4-table1.

### Public databases and analysis methods

Transcriptome sequencing data of 142 TNBC and 695 non-TNBC clinical samples were selected from TCGA database (https://www.cancer.gov/ccg/research/genome-sequencing/tcga), differential gene analysis was performed with the R package ‘limma’. IGF2BP3 and c-Met protein expression data were extracted from CPTAC database (https://proteomics.cancer.gov/programs/cptac), gene correlation analysis was performed with the R package ‘corrplot’.

### Breast cancer samples

TNBC breast cancer tissue samples were collected from Jiangsu Breast Disease Center, the First Affiliated Hospital with Nanjing Medical University. All samples were not received preoperative neoadjuvant chemotherapy and were diagnosed as TNBC in post-operative immunohistochemistry. The use of breast cancer samples was approved by the Institutional Ethics Committee of the First Affiliated Hospital of Nanjing Medical University.

### Lung metastasis of breast cancer model in mice

The balb/c female nude mice were from animal research center of Nanjing Medical University (Nanjing, China). 1.5×10^6^ cells transfected with NC+Vector, NC+c-Met, shIGF2BP3+Vector, shIGF2BP3+c-Met were injected into the nude mice through the tail vein. After 4 weeks, the lungs were sold out and photographed, fixed with 4 % paraformaldehydeand, embedded in paraffin, sectioned for hematoxylin-eosin staining and immunohistochemistry staining afterwards. All animal experiments were approved by Institutional Animal Care and Use Committee of Nanjing Medical University.

### Statistical analysis

All data were analyzed and presented by GraphPad Prism (Version 8.0). Student t-test and one-way ANOVA were used for comparing differences between groups. All experiments were repeated independently for three times, error bars represent SEM.*\*P*<0.05 were deemed to statistically significant.

## Acknowledgments

We would like to thank JW from department of pathology for technical instruction.

The pattern diagram is drawn with https://www.Figuredraw.com.

## Data Availability

The data generated in this study can be obtained from the corresponding author upon reasonable request.

## References

Abd El-Aziz, Y. S., Gillson, J., Jansson, P. J., & Sahni, S. (2022). Autophagy: A promising target for triple negative breast cancers. Pharmacological Research, 175, 106006. doi:10.1016/j.phrs.2021.106006

An, Y., & Duan, H. (2022). The role of m6A RNA methylation in cancer metabolism. Molecular Cancer, 21(1), 14. doi:10.1186/s12943-022-01500-4

Bai, X., Ni, J., Beretov, J., Graham, P., & Li, Y. (2021). Triple-negative breast cancer therapeutic resistance: Where is the Achilles’ heel? Cancer Letters, 497, 100–111. doi:10.1016/j.canlet.2020.10.016

Bianchini, G., Balko, J. M., Mayer, I. A., Sanders, M. E., & Gianni, L. (2016). Triple-negative breast cancer: challenges and opportunities of a heterogeneous disease. Nature Reviews. Clinical Oncology, 13(11), 674–690. doi:10.1038/nrclinonc.2016.66

Bianchini, G., De Angelis, C., Licata, L., & Gianni, L. (2022). Treatment landscape of triple-negative breast cancer - expanded options, evolving needs. Nature Reviews. Clinical Oncology, 19(2). doi:10.1038/s41571-021-00565-2

Brito Querido, J., Díaz-López, I., & Ramakrishnan, V. (2024). The molecular basis of translation initiation and its regulation in eukaryotes. Nature Reviews. Molecular Cell Biology, 25(3), 168–186. doi:10.1038/s41580-023-00624-9

Chen, H.-T., Liu, H., Mao, M.-J., Tan, Y., Mo, X.-Q., Meng, X.-J.,… Jiang, G.-M. (2019). Crosstalk between autophagy and epithelial-mesenchymal transition and its application in cancer therapy. Molecular Cancer, 18(1), 101. doi:10.1186/s12943-019-1030-2

Chen, X., Gong, W., Shao, X., Shi, T., Zhang, L., Dong, J.,… Guo, B. (2022). METTL3-mediated m6A modification of ATG7 regulates autophagy-GATA4 axis to promote cellular senescence and osteoarthritis progression. Annals of the Rheumatic Diseases, 81(1), 87–99. doi:10.1136/annrheumdis-2021-221091

Dower, C. M., Wills, C. A., Frisch, S. M., & Wang, H.-G. (2018). Mechanisms and context underlying the role of autophagy in cancer metastasis. Autophagy, 14(7), 1110–1128. doi:10.1080/15548627.2018.1450020

Gerstberger, S., Jiang, Q., & Ganesh, K. (2023). Metastasis. Cell, 186(8), 1564–1579. doi:10.1016/j.cell.2023.03.003

Gundamaraju, R., Lu, W., Paul, M. K., Jha, N. K., Gupta, P. K., Ojha, S.,… Ghavami, S. (2022). Autophagy and EMT in cancer and metastasis: Who controls whom? Biochimica Et Biophysica Acta. Molecular Basis of Disease, 1868(9), 166431. doi:10.1016/j.bbadis.2022.166431

Hao, W., Dian, M., Zhou, Y., Zhong, Q., Pang, W., Li, Z.,… Xiao, D. (2022). Autophagy induction promoted by m6A reader YTHDF3 through translation upregulation of FOXO3 mRNA. Nature Communications, 13(1), 5845. doi:10.1038/s41467-022-32963-0

He, M., Lei, H., He, X., Liu, Y., Wang, A., Ren, Z.,… Yang, L. (2022). METTL14 Regulates Osteogenesis of Bone Marrow Mesenchymal Stem Cells via Inducing Autophagy Through m6A/IGF2BPs/Beclin-1 Signal Axis. Stem Cells Translational Medicine, 11(9). doi:10.1093/stcltm/szac049

Hu, M., Chen, J., Liu, S., & Xu, H. (2023). The Acid Gate in the Lysosome. Autophagy, 19(4), 1368–1370. doi:10.1080/15548627.2022.2125629

Huang, H., Weng, H., & Chen, J. (2020). m6A Modification in Coding and Non-coding RNAs: Roles and Therapeutic Implications in Cancer. Cancer Cell, 37(3), 270–288. doi:10.1016/j.ccell.2020.02.004

Huang, H., Weng, H., Sun, W., Qin, X., Shi, H., Wu, H.,… Chen, J. (2018). Recognition of RNA N6-methyladenosine by IGF2BP proteins enhances mRNA stability and translation. Nature Cell Biology, 20(3), 285–295. doi:10.1038/s41556-018-0045-z

Jackson, R. J., Hellen, C. U. T., & Pestova, T. V. (2010). The mechanism of eukaryotic translation initiation and principles of its regulation. Nature Reviews. Molecular Cell Biology, 11(2), 113–127. doi:10.1038/nrm2838

Jiang, T., He, X., Zhao, Z., Zhang, X., Wang, T., & Jia, L. (2022). RNA m6A reader IGF2BP3 promotes metastasis of triple-negative breast cancer via SLIT2 repression. FASEB Journal : Official Publication of the Federation of American Societies For Experimental Biology, 36(11), e22618. doi:10.1096/fj.202200751RR

Kim, H.-Y., Ha Thi, H. T., & Hong, S. (2018). IMP2 and IMP3 cooperate to promote the metastasis of triple-negative breast cancer through destabilization of progesterone receptor. Cancer Letters, 415, 30–39. doi:10.1016/j.canlet.2017.11.039

Kim, Y. C., & Guan, K.-L. (2015). mTOR: a pharmacologic target for autophagy regulation. The Journal of Clinical Investigation, 125(1), 25–32. doi:10.1172/JCI73939

Li, J., Kim, S. G., & Blenis, J. (2014). Rapamycin: one drug, many effects. Cell Metabolism, 19(3), 373–379. doi:10.1016/j.cmet.2014.01.001

Liang, D., Lin, W.-J., Ren, M., Qiu, J., Yang, C., Wang, X.,… Wang, W. (2022). m6A reader YTHDC1 modulates autophagy by targeting SQSTM1 in diabetic skin. Autophagy, 18(6), 1318–1337. doi:10.1080/15548627.2021.1974175

Liberman, N., Gandin, V., Svitkin, Y. V., David, M., Virgili, G., Jaramillo, M.,… Sonenberg, N. (2015). DAP5 associates with eIF2β and eIF4AI to promote Internal Ribosome Entry Site driven translation. Nucleic Acids Research, 43(7), 3764–3775. doi:10.1093/nar/gkv205

Lin, S., Choe, J., Du, P., Triboulet, R., & Gregory, R. I. (2016). The m(6)A Methyltransferase METTL3 Promotes Translation in Human Cancer Cells. Molecular Cell, 62(3), 335–345. doi:10.1016/j.molcel.2016.03.021

Lin, X., Chai, G., Wu, Y., Li, J., Chen, F., Liu, J.,… Wang, H. (2019). RNA m6A methylation regulates the epithelial mesenchymal transition of cancer cells and translation of Snail. Nature Communications, 10(1), 2065. doi:10.1038/s41467-019-09865-9

Lin, Z., Li, J., Zhang, J., Feng, W., Lu, J., Ma, X.,… Zhang, X. (2023). Metabolic Reprogramming Driven by IGF2BP3 Promotes Acquired Resistance to EGFR Inhibitors in Non-Small Cell Lung Cancer. Cancer Research, 83(13), 2187–2207. doi:10.1158/0008-5472.CAN-22-3059

Liu, L., Li, H., Hu, D., Wang, Y., Shao, W., Zhong, J.,… Zhang, J. (2022). Insights into N6-methyladenosine and programmed cell death in cancer. Molecular Cancer, 21(1), 32. doi:10.1186/s12943-022-01508-w

Massagué, J., & Ganesh, K. (2021). Metastasis-Initiating Cells and Ecosystems. Cancer Discovery, 11(4), 971–994. doi:10.1158/2159-8290.CD-21-0010

Meyer, K. D., Patil, D. P., Zhou, J., Zinoviev, A., Skabkin, M. A., Elemento, O.,… Jaffrey, S. R. (2015). 5’ UTR m(6)A Promotes Cap-Independent Translation. Cell, 163(4). doi:10.1016/j.cell.2015.10.012

Nag, S., Goswami, B., Das Mandal, S., & Ray, P. S. (2022). Cooperation and competition by RNA-binding proteins in cancer. Seminars In Cancer Biology, 86(Pt 3), 286–297. doi:10.1016/j.semcancer.2022.02.023

Ning, J., Pei, Z., Wang, M., Hu, H., Chen, M., Liu, Q.,… Zhang, R. (2023). Site-specific Atg13 methylation-mediated autophagy regulates epithelial inflammation in PM2.5-induced pulmonary fibrosis. Journal of Hazardous Materials, 457, 131791. doi:10.1016/j.jhazmat.2023.131791

Nolan, E., Lindeman, G. J., & Visvader, J. E. (2023). Deciphering breast cancer: from biology to the clinic. Cell, 186(8), 1708–1728. doi:10.1016/j.cell.2023.01.040

Singla, M., & Bhattacharyya, S. (2017). Autophagy as a potential therapeutic target during epithelial to mesenchymal transition in renal cell carcinoma: An in vitro study. Biomedicine & Pharmacotherapy = Biomedecine & Pharmacotherapie, 94, 332–340. doi:10.1016/j.biopha.2017.07.070

So, L., Lee, J., Palafox, M., Mallya, S., Woxland, C. G., Arguello, M.,… Fruman, D. A. (2016). The 4E-BP-eIF4E axis promotes rapamycin-sensitive growth and proliferation in lymphocytes. Science Signaling, 9(430), ra57. doi:10.1126/scisignal.aad8463

Su, Z., Yang, Z., Xu, Y., Chen, Y., & Yu, Q. (2015). Apoptosis, autophagy, necroptosis, and cancer metastasis. Molecular Cancer, 14, 48. doi:10.1186/s12943-015-0321-5

Thiery, J. P., Acloque, H., Huang, R. Y. J., & Nieto, M. A. (2009). Epithelial-mesenchymal transitions in development and disease. Cell, 139(5), 871–890. doi:10.1016/j.cell.2009.11.007

Wang, A., Huang, H., Shi, J.-H., Yu, X., Ding, R., Zhang, Y.,… Zou, Q. (2023). USP47 inhibits m6A-dependent c-Myc translation to maintain regulatory T cell metabolic and functional homeostasis. The Journal of Clinical Investigation, 133(23). doi:10.1172/JCI169365

Wang, H., Zhang, Y., Wu, Q., Wang, Y.-B., & Wang, W. (2018). miR-16 mimics inhibit TGF-β1-induced epithelial-to-mesenchymal transition via activation of autophagy in non-small cell lung carcinoma cells. Oncology Reports, 39(1), 247–254. doi:10.3892/or.2017.6088

Wang, X., Tian, L., Li, Y., Wang, J., Yan, B., Yang, L.,… Sun, Y. (2021). RBM15 facilitates laryngeal squamous cell carcinoma progression by regulating TMBIM6 stability through IGF2BP3 dependent. Journal of Experimental & Clinical Cancer Research : CR, 40(1), 80. doi:10.1186/s13046-021-01871-4

Wang, X., Zhao, B. S., Roundtree, I. A., Lu, Z., Han, D., Ma, H.,… He, C. (2015). N(6)-methyladenosine Modulates Messenger RNA Translation Efficiency. Cell, 161(6), 1388–1399. doi:10.1016/j.cell.2015.05.014

Yang, Y., Li, Q., Ling, Y., Leng, L., Ma, Y., Xue, L.,… Tao, S. (2022). m6A eraser FTO modulates autophagy by targeting SQSTM1/P62 in the prevention of canagliflozin against renal fibrosis. Frontiers In Immunology, 13, 1094556. doi:10.3389/fimmu.2022.1094556

Zhang, N., Shen, Y., Li, H., Chen, Y., Zhang, P., Lou, S., & Deng, J. (2022). The m6A reader IGF2BP3 promotes acute myeloid leukemia progression by enhancing RCC2 stability. Experimental & Molecular Medicine, 54(2), 194–205. doi:10.1038/s12276-022-00735-x

Zhang, X., Shi, L., Sun, H.-D., Wang, Z.-W., Xu, F., Wei, J.-F., & Ding, Q. (2023). IGF2BP3 mediates the mRNA degradation of NF1 to promote triple-negative breast cancer progression via an m6A-dependent manner. Clinical and Translational Medicine, 13(9), e1427. doi:10.1002/ctm2.1427

Zhang, Y., Xia, M., Jin, K., Wang, S., Wei, H., Fan, C.,… Xiong, W. (2018). Function of the c-Met receptor tyrosine kinase in carcinogenesis and associated therapeutic opportunities. Molecular Cancer, 17(1), 45. doi:10.1186/s12943-018-0796-y

Zhao, B. S., Roundtree, I. A., & He, C. (2017). Post-transcriptional gene regulation by mRNA modifications. Nature Reviews. Molecular Cell Biology, 18(1), 31–42. doi:10.1038/nrm.2016.132

Zhu, T.-Y., Hong, L.-L., & Ling, Z.-Q. (2023). Oncofetal protein IGF2BPs in human cancer: functions, mechanisms and therapeutic potential. Biomarker Research, 11(1), 62. doi:10.1186/s40364-023-00499-0

